## Supplementary_Information for "“Lose-to-gain” adaptation to genome decay in the structure of the smallest eukaryotic ribosomes"

**Affiliations**

### To whom correspondence should be addressed:

**Key words:**

**Supplementary figures**

Figure S1 | Collection and processing of cryo-EM data.

Figure S2 | Overall and local resolution assessment of the *E. cuniculi* ribosome maps.

Figure S3 | Additional truncations in rRNA and protein segments in *E. cuniculi* ribosomes compared to *Nosematedae* ribosomes from *P. locustae* and *V. necatrix*.

Figure S4 | Progressive degeneration of a ribosomal protein upon transition from free-living fungi with large genomes to microsporidians with extremely reduced genomes.

Figure S5 | Local changes in ribosomal protein structure compensate for rRNA reduction.

Figure S6 | A spermidine-like small molecule at the protein-protein interface in the interior of the *E. cuniculi* ribosome.

**Supplementary tables**

Table S1 | Validation of the *E. cuniculi* ribosome structure.

**Supplementary data** (uploaded as separate files)

Supplementary Data S1 | A script to retrieve eL20 sequences from UniProt.

Supplementary Data S2 | Eukaryotic eL20 sequences analyzed in this study (in this file, sequences are named by the name of a corresponding organism and the number of protein-coding genes in this organism)

Supplementary Data S3 | Cryo-EM maps and the structure produced in this study:

- The large ribosomal subunit: 2021_Ecuniculi_ribosome_LSU.mrc
- The head of the small subunit: 2021_Ecuniculi_ribosome_SSU_head.mrc
- The body of the small subunit: 2021_Ecuniculi_ribosome_SSU_body.mrc
- The 80S ribosome from E. cuniculi: 2021_Ecuniculi_ribosome.pdb

Please download using this link to download maps:
https://drive.google.com/drive/folders/1Sq3ufbPkJjdiD3AMwmJe6hlSbzlIjrN7?usp=sharing

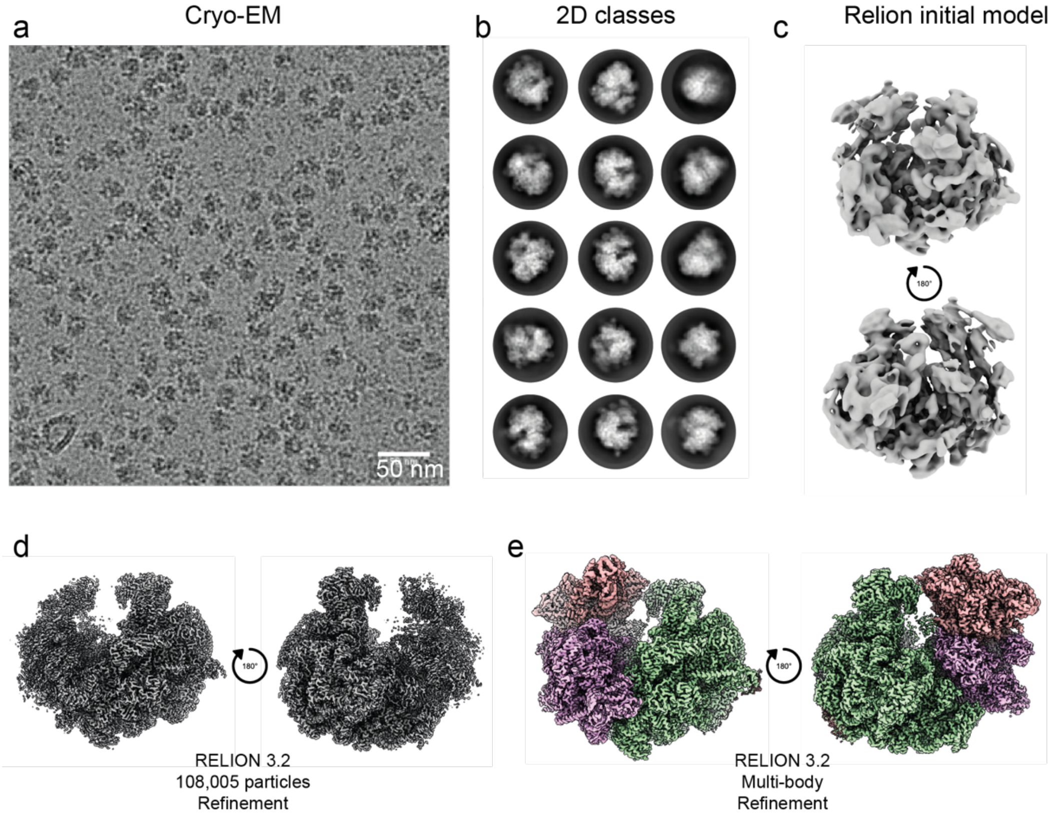


**Figure S1 | Collection and processing of cryo-EM data.** (**a**) and Representative cryo-electron microscopy micrograph of *E. cuniculi* ribosomes (**b**) Representative cryo-EM 2D class averages (**c**) and initial model generated in RELION 3.1. Cryo-EM density following (**d**) 3D refinement and (**e**) multi-body refinement. Final refinement was performed on 108,005 particles extracted from 2,210 micrographs in RELION 3.1. The 3 bodies "LSU", "SSU-head" and "SSU-body" masks were used for multibody refinement.

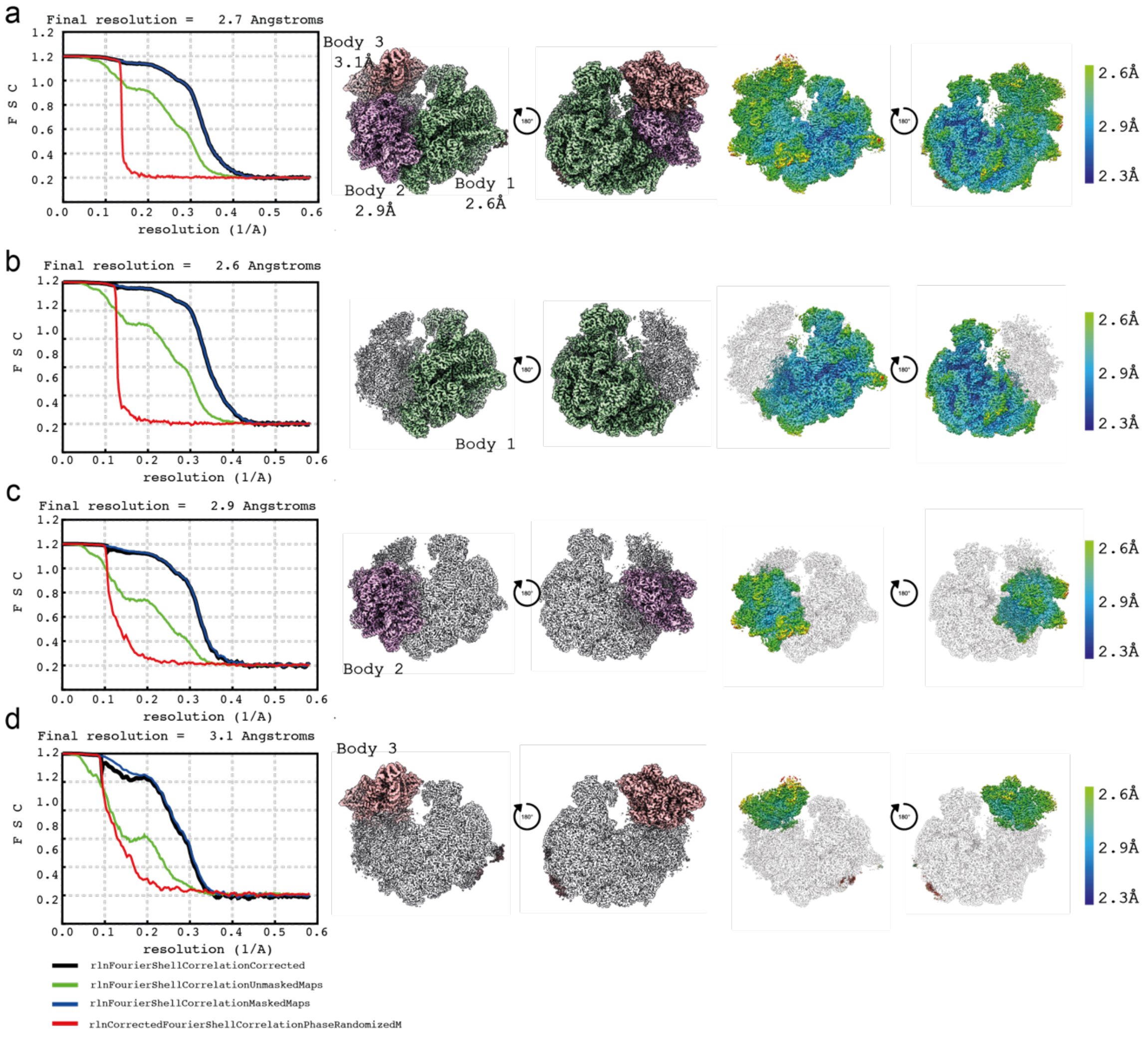


**Figure S2 | Overall and local resolution assessment of the *E. cuniculi* ribosome maps.** (**a-d**) Overall and local resolution estimations of (**a**) the consensus map, (**b**) the LSU map, (c) the SSU-body map, (**d**) the SSU-head map. The panel on the left shows Fourier Shell Correlation (FSC) curves for the respective map. An overview of the cryo-EM map is shown next to a slab view and two 180-degree related views colored according to local resolution. Local resolution was calculated using RELION 3.1.

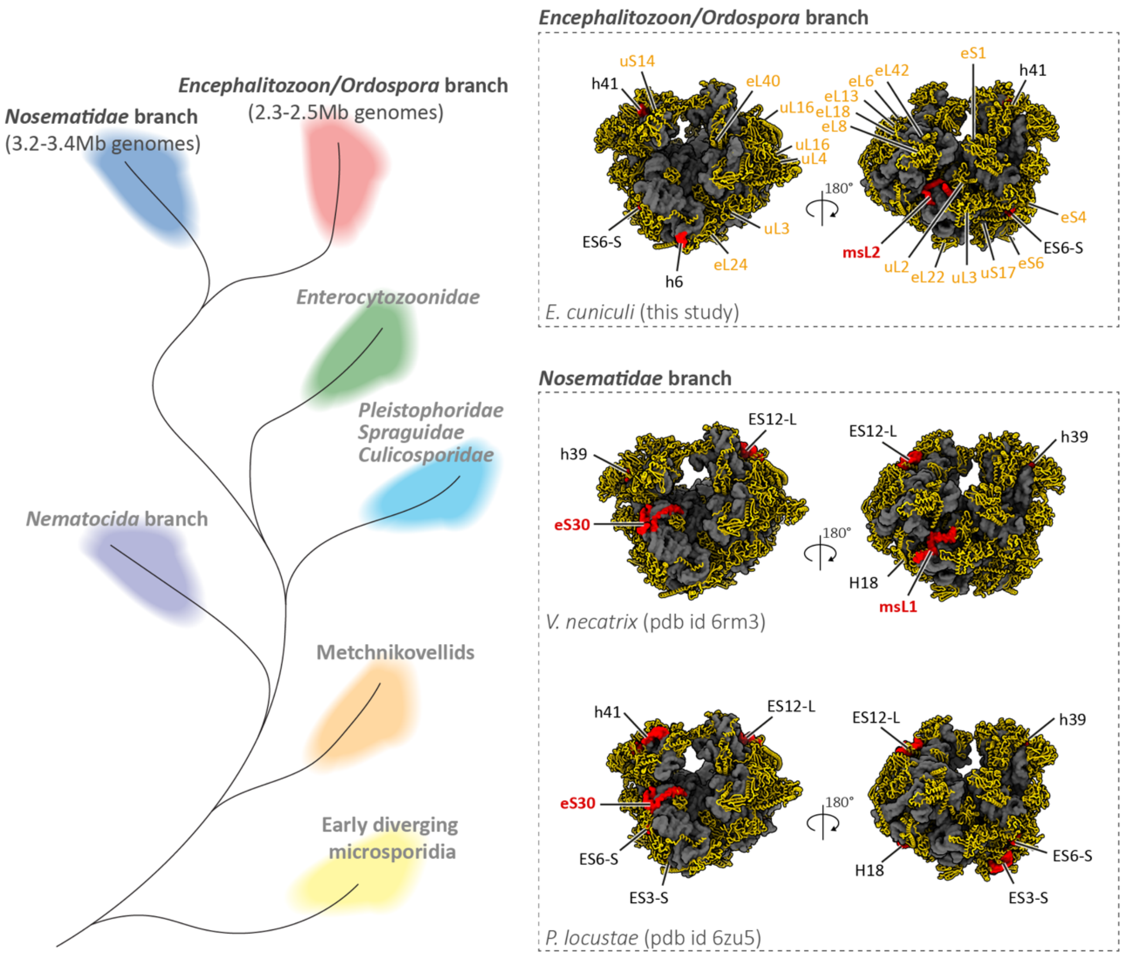


**Figure S3** | *E. cuniculi* ribosomes possess additional truncations in rRNA and protein segments compared to *Nosematedae* ribosomes from *P. locustae* and *V. necatrix*. Microsporidian tree of life is shown to illustrate *Encephalitozoon/Ordospora* branch of microsporidian species, which are known for their smallest and most reduced genomes among eukaryotic species.

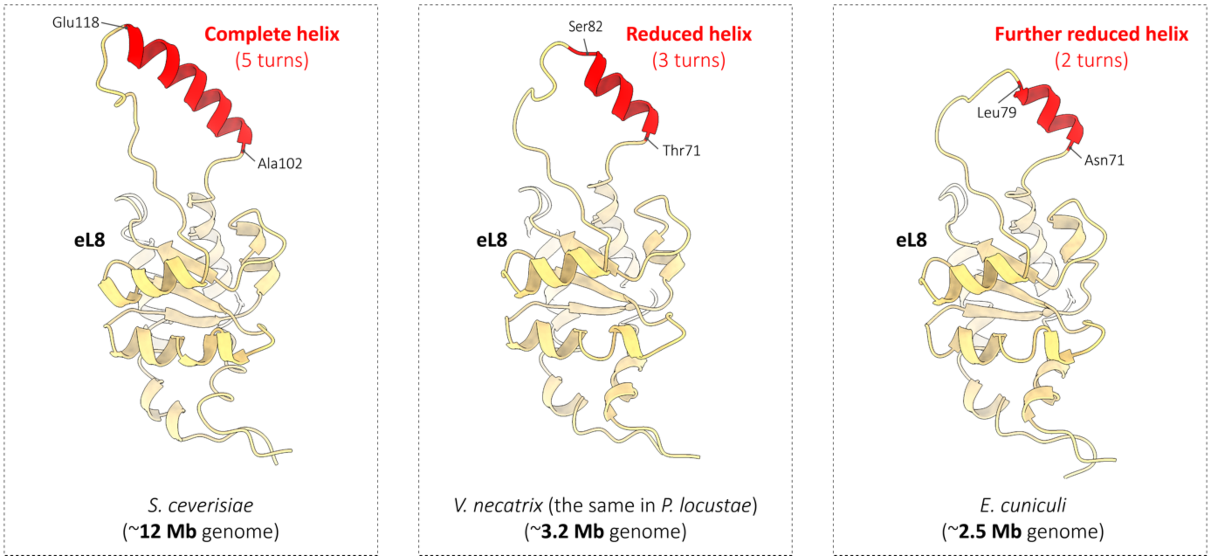


**Figure S4** | Progressive degeneration of a ribosomal protein upon transition from free-living fungi with large genomes to microsporidians with extremely reduced genomes.

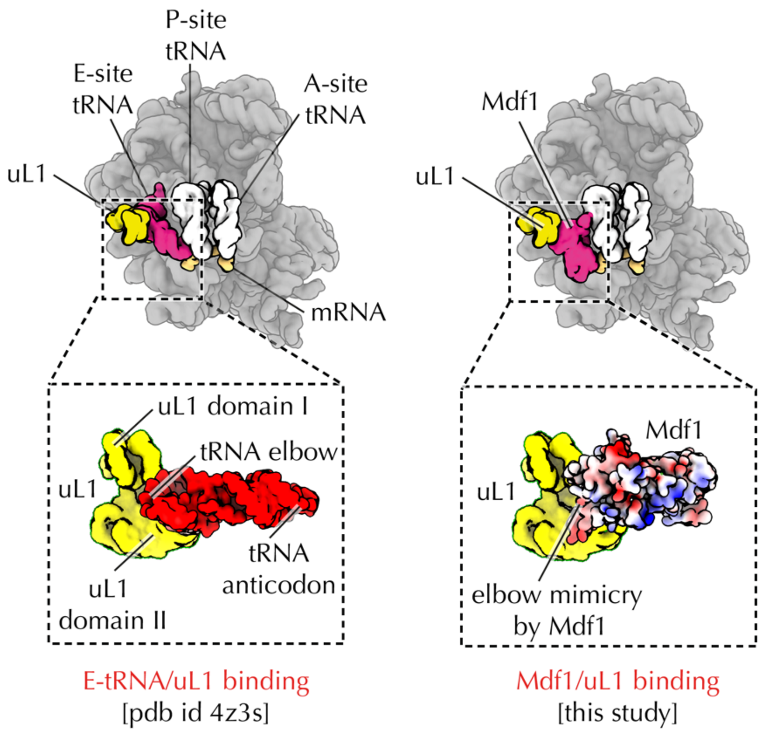


**Fig S5 | Ribosome hibernation factor Mdf1 mimics deacetylated tRNAs.** Mdf1 was initially described in microsporidian parasites *V. necatrix*. Similar to other hibernation factors, Mdf1 was shown to block a ribosomal E site helping inactivate ribosomes when parasites sporulate and become metabolically inactive. Our structure revealed that Mdf1 not only binds *E. cuniculi* ribosomes at the E site but also forms a previously unknown contact with the ribosomal L1-stalk (the part of the ribosome that helps release deacylated tRNAs from the ribosome during protein synthesis) (**Fig. 6**). In so doing, Mdf1 exploits the same contacts as the "elbow"-segment of deacylated tRNA molecules. This previously unknown molecular mimicry suggests that Mdf1 dissociates from the ribosome using the same mechanism as deacetylated tRNAs, explaining how ribosomes can remove this hibernation factor to reactivate protein synthesis.

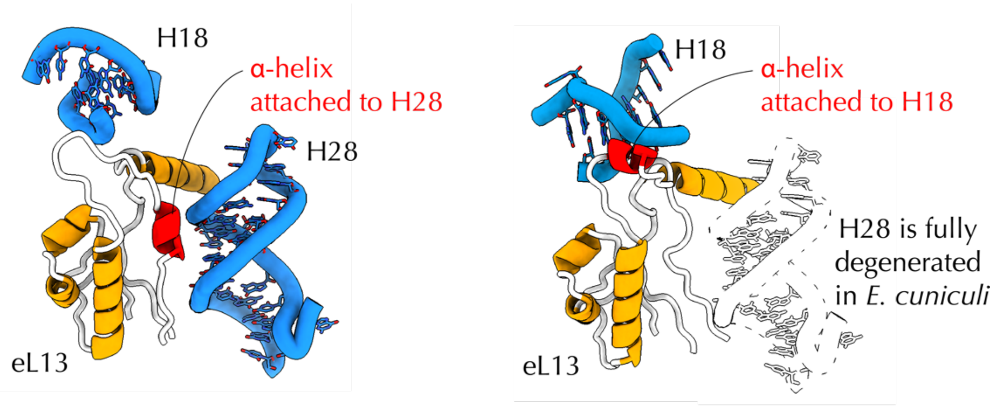


**Figure S6 | Microsporidian ribosomal proteins reinvent their rRNA attachment sites as an apparent adaption to rRNA degeneration.**

Previously, we described a mechanism by which ribosomes adapt to the evolution of new rRNA segments in eukaryotic ribosomes. We showed that eukaryotic proteins (compared to their bacterial homologs) remodeled their solvent-exposed loops into miniature $\alpha$-helices. These helices serve as new binding sites for eukaryote-specific rRNA expansions, ensuring a proper folding of rRNA expansions in the ribosome.

Here, we found that the same strategy is used by *E. cuniculi* ribosomes to help adapt ribosomes to rRNA degeneration. For example, protein eL13 converts one of its loops into an $\alpha$-helix that binds the truncated rRNA. This structural change allows eL13 to reinforce its binding to the *E. cuniculi* ribosome in which the primary binding site of eL13 is eliminated due to the loss the rRNA expansion $ES7^{L}$ (and the helix H28).

This finding shows that, by transforming their structure back and forth between the states of "loop" and "$\alpha$-helix", ribosomal proteins can create and eliminate rRNA binding sites, thus heling rRNA to expand or shrink without loss of the ribosome integrity. This strategy may explain numerous changes in ribosomal proteins from other species that possess anomalously long or short rRNA molecules.

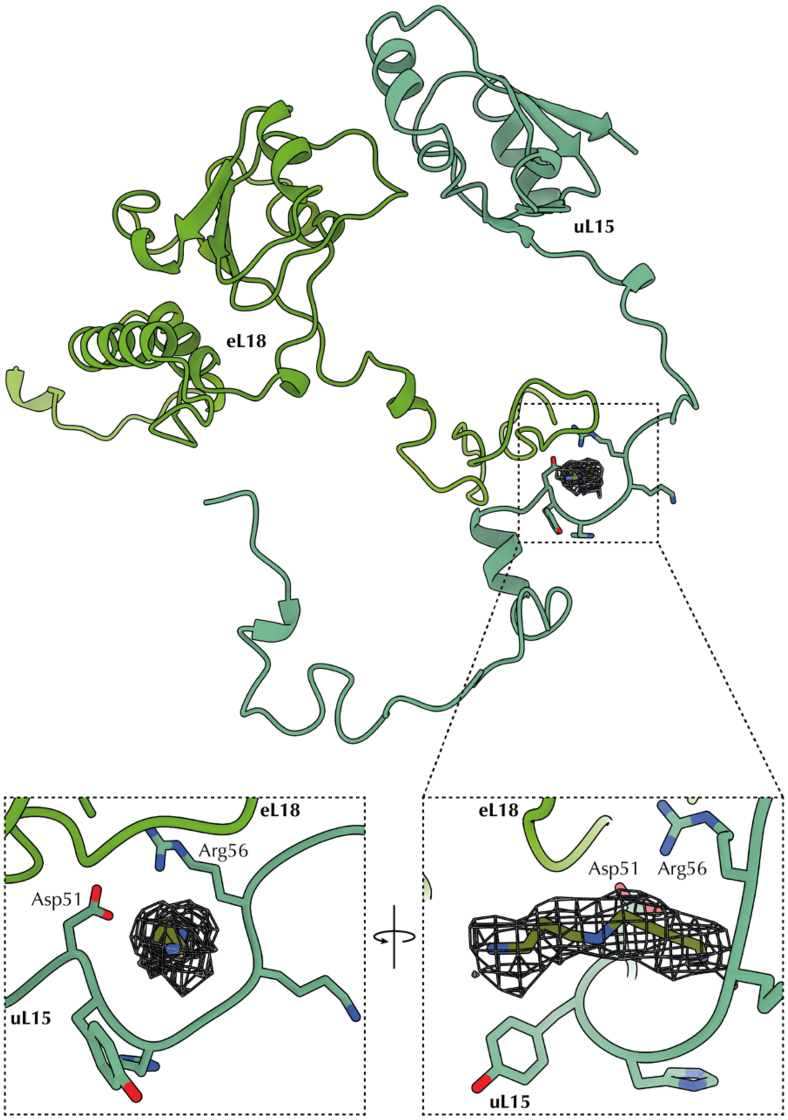


**Figure S7 | A spermidine-like small molecule trapped at a protein-protein interface in the interior of the *E. cuniculi* ribosome.** A fragment of the cryo-EM structure shows the cryo-EM map (contoured at 0.1V) of a small molecule that is trapped in the ribosome interior between ribosomal proteins uL15 and eL18. A salt-bridge between the residues Asp51 and Arg56 in uL15 creates a donut-like binding site for this molecule in the ribosome structure. In other eukaryotic ribosomes (e.g. in yeasts and humans), this void is occupied by the C-terminal extension of protein eL18, which is truncated in *E. cuniculi*.

| **Model composition** | | **R.m.s. deviations**[a] | |
|---|---|---|---|
| Non-hydrogen atoms[a] | 166129 | Bond lengths (Å) | 0.009 |
| Protein residues[b] | 10885 | Bond angles (°) | 0.943 |
| Nucleic acid residues[b] | 3734 | **Protein geometry validation**[a] | |
| Metal ions[a] | 8 Zn | Rotamer outliers (%) | 0.09 |
| Ligands | spermidine AMP | Ramachandran outliers (%) | 0.21 |
| **General validation**[a] | | Ramachandran favoured (%) | 90.10 |
| CC (model to map fit) [c] | 0.77 | **RNA geometry validation**[b] | |
| Clashscore | 8.35 | Sugar pucker outliers (%) | 0.80 |
| MolProbity score | 2.00 | Backbone conformation outlie (%) | 20.30 |

**Table S1 Model validation statistics.** [a]Obtained from Phenix refine log and phenix molprobity.
[b]Obtained from MolProbity web server.
[c]CC = correlation coefficient, measure of fit into consensus map.
